# Enhanced Cell Tracking Using A GAN-based Super-Resolution Video-to-Video Time-Lapse Microscopy Generative Model

**DOI:** 10.1101/2024.06.11.598572

**Authors:** Abolfazl Zargari, Najmeh Mashhadi, S. Ali Shariati

## Abstract

Cells are among the most dynamic entities, constantly undergoing various processes such as growth, division, movement, and interaction with other cells as well as the environment. Time-lapse microscopy is central to capturing these dynamic behaviors, providing detailed temporal and spatial information that allows biologists to observe and analyze cellular activities in real-time. The analysis of time-lapse microscopy data relies on two fundamental tasks: cell segmentation and cell tracking. Integrating deep learning into bioimage analysis has revolutionized cell segmentation, producing models with high precision across a wide range of biological images. However, developing generalizable deep-learning models for tracking cells over time remains challenging due to the scarcity of large, diverse annotated datasets of time-lapse movies of cells. To address this bottleneck, we propose a GAN-based time-lapse microscopy generator, termed tGAN, designed to significantly enhance the quality and diversity of synthetic annotated time-lapse microscopy data. Our model features a dual-resolution architecture that adeptly synthesizes both low and high-resolution images, uniquely capturing the intricate dynamics of cellular processes essential for accurate tracking. We demonstrate the performance of tGAN in generating high-quality, realistic, annotated time-lapse videos. Our findings indicate that tGAN decreases dependency on extensive manual annotation to enhance the precision of cell tracking models for time-lapse microscopy.

## Introduction

Time-lapse microscopy, a crucial tool in cell biology, captures the dynamics of cellular processes with high temporal resolution at single cell level. However, the analysis of time-lapse images poses significant challenges, particularly in tracking cells over time [1]. A key hurdle in this process is the lack of annotated datasets, which are vital for training deep learning models to accurately segment and track individual cells over time [2]. Annotated time-lapse microscopy images are indispensable for understanding cellular mechanisms and responses, yet creating such datasets is often labor-intensive and requires expert knowledge.

Recent advancements in deep learning, particularly in generative adversarial networks (GANs) [3], offer an unprecedented opportunity for the generation of realistic synthetic data [4,5]. GAN models, known for their ability to generate highly realistic synthetic data, have found applications across a wide range of disciplines, from art creation to medical imaging [6-8]. GAN models provide innovative solutions to simulate realistic biological data, which is essential for training and testing analytical models [9]. Among these applications, the generation of synthetic cell images from time-lapse microscopy using deep learning models, especially GANs, stands out as a particularly promising area [10]. These models offer new avenues for research in cell biology, enabling the creation of detailed and accurate representations of cellular processes.

To address the paucity of annotated time-lapse microscopy images, we introduce a video-to-video generative approach designed to generate annotated time-lapse microscopy images. Our time-lapse generative model, termed tGAN, converts binary mask image sequences into corresponding high-resolution, synthetic images of cells. In addition to the generation of realistic time-lapse videos of cells, tGAN can be used to train tracking models with limited annotated datasets. The model offers a significant advancement in the automated generation of annotated time-lapse microscopy datasets, facilitating more efficient and accurate analysis of cellular dynamics.

## Results

### Designing a novel GAN-based super-resolution video-to-video generator

The architectural design of our video-to-video generative model is central to its performance. Our design is characterized by its two-part structure, encompassing both video-to-video low-resolution and image-to-image super-resolution models (Figure 1A). While the low-resolution model adequately captures the dynamics of time-lapse microscopy videos, our independent super-resolution model further enhances the quality of synthetic images by adding morphological details to single cells. This two-part structure is designed to optimize computational efficiency and precision. Since the low-resolution model training process involves multiple subunits such as discriminators, flow networks, and the video-to-video generator (Figure 1B), integrating all these elements at a high resolution would significantly increase computational costs and could potentially degrade the performance and training efficiency of the generator. Therefore, we initially trained the video-to-video generator in low resolution, achieving higher accuracy with less computational overhead. Subsequently, we trained a GAN-based super-resolution model (Figure 1C) to refine the video-to-video model outputs to high resolution, enhancing the quality and adding additional morphological details to cells for applications requiring high fidelity. This sequential approach not only ensures detailed, high-quality outputs but also maintains manageable computational demands, facilitating more efficient processing and superior performance in scenarios that require detailed, high-resolution images.

**Figure 1.**
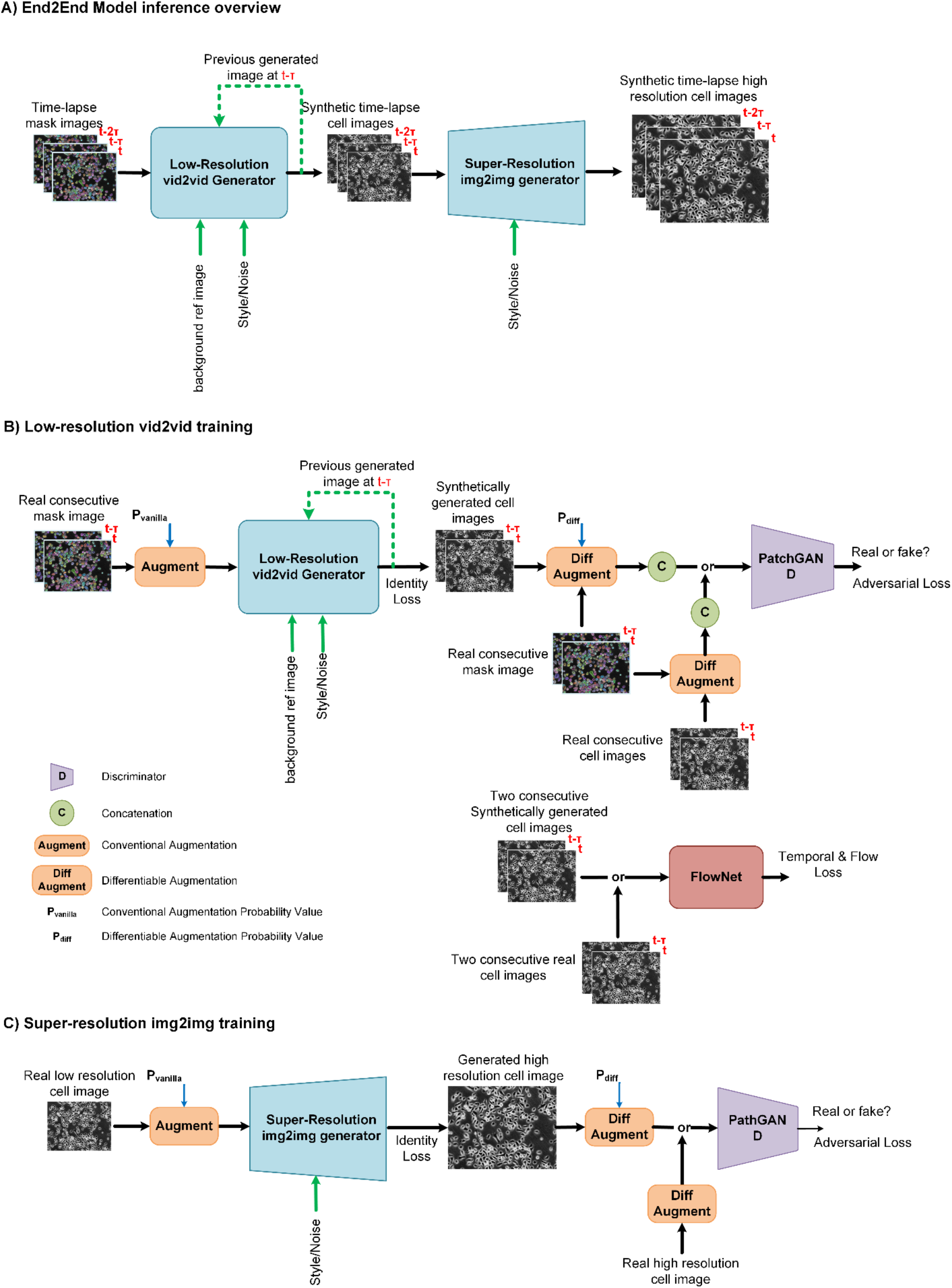
Overview of the proposed GAN-based generator. A) Illustration of the two-part inference architecture: This includes both the low-resolution video-to-video and super-resolution image-to-image models, where the initial phase generates low-resolution time-lapse microscopy image sequences to capture essential cellular dynamics, followed by the super-resolution phase that refines these sequences into more detailed and high-quality images. B) Low-resolution video-to-video training process: Showcasing the sequence-based approach and the integration of various inputs and deep learning components. C) Super-resolution image-to-image training process: Highlighting the approach and models used for refining textural details and enhancing aesthetic elements, thereby producing scientifically accurate and visually high-resolution images.

The low-resolution video-to-video generator, inspired by 2D-UNET architecture [11], accepts multiple inputs, including current n (for example, two) consecutive mask frames, a previously generated cell frame (at t-T), and a consistent reference background image, ensuring context-aware and consistent background generation (Figure 1B and Figure S2). One of the distinctive features of this phase is the inclusion of a reference background image. Incorporating a reference background image in our low-resolution GAN model enhances contextual accuracy to ensure consistency in background features. By increasing the variability of background visual features in synthetic samples, tGAN can accommodate various real-world scenarios like variable background noises, debris, or optical artifacts common in microscopy. This approach enriches the realism and detail of the generated images, which is crucial for precise analysis in applications such as time-lapse microscopy, where background features can change over time. The integration of attention layers allows for adaptively integrating visual features from three model inputs. The inclusion of style and noise injection into the decoding path adds variability and realism to the generated images, as we have validated in our prior research [12],

Transitioning to the super-resolution phase, our image-to-image generator employs an enhanced UNET-based architecture, incorporating style and noise injection for refining textural details and aesthetic elements, adding finer details and thus resulting in higher quality and more detailed images (Figure 1C and Figure S3). While, in our experiments, we targeted a high resolution of 512×768 for the super-resolution model output, our model is adaptable and can be easily configured to achieve higher resolutions by adjusting its hyperparameters. The discriminators, designed with a PatchGAN architecture [13] and enhanced with a linear attention layer [14], effectively distinguish fine details in images (Figure 1B and Figure S4). This enhances the model’s accuracy in differentiating between real and synthetic images. In the training process of the video-to-video generator, alongside the GAN and discriminator models, we concurrently trained and used a FlowNET (Figure 1B and Figure S5) designed to calculate and integrate flow loss into the training regime. This FlowNET plays a crucial role in determining the optical flow loss [15], comparing motion between consecutive frames in both real and generated sequences. This is essential for preserving the dynamic nature of cell movements in time-lapse microscopy. The optical flow loss computed by our FlowNet ensures that the temporal coherence and motion consistency of generated images align closely with actual microscopy sequences. However, it is not incorporated in the super-resolution model, which concentrates on image-to-image translation rather than the generation of video sequences, thereby making flow consistency less relevant in that context.

We selected loss functions, including temporal consistency and perceptual losses, that ensure temporal coherence and visual similarity between generated and real images (Equations 1-3). Throughout the training process, we also employed augmentation strategies, including video-level augmentations for general model training, as well as video-level differentiable augmentations [16] for the discriminators (Figure S6). These augmentations enhance the model’s robustness and prevent overfitting, especially when the training dataset size is limited, as we have shown previously [12].

### Benchmarking tGAN performance for generation of annotated video frames

To evaluate the performance of our model, we compared our model with the vid2vid model, a notable example among state-of-the-art video-to-video generative models (Table 1) [17]. The vid2vid framework is particularly relevant for our comparative analysis as it is among very few publicly available generative models capable of converting mask scene sequences into realistic scene sequences and generating high-resolution video frames. We used the Structural Similarity Index (SSIM) and Peak Signal-to-Noise Ratio (PSNR) to compare image quality and similarity to real frames between the vid2vid and the models [18]. Additionally, we utilized the Frechet Video Distance (FVD) [19] to measure the distributional similarity between generated and real images, providing insights into the perceptual quality of tGAN’s outputs. To evaluate the temporal coherence of generated video sequences, we adopted specific metrics designed to assess the smoothness and continuity of changes across frames. Finally, the Perceptual Image Patch Similarity (LPIPS) metric [20] was used to gauge the perceptual resemblance of generated images to real ones, ensuring that tGAN’s outputs align closely with human visual judgment. Together, these metrics provided a robust framework for evaluating and benchmarking our model for video-to-video generation tasks. As presented in Table 1, our tGAN model obtained better performance across almost all the metrics and five cell types when tested on the DeepSea [21] and Cell Tracking Challenge [22] dataset time-lapse image sequences. Figure 2 showcases examples of two consecutive frames generated by our tGAN for each of the three DeepSea cell types, which are compared with the corresponding outputs from the vid2vid model. tGAN-generated images successfully capture realistic details of the cell bodies, including features like the nucleus, as well as the nuances of the background, demonstrating the model’s effectiveness in rendering intricate biological structures. Figure S1 also compares the performance of our tGAN output with the vid2vid model, specifically focusing on the ability to replicate fine visual features of single stem cells. This comparison highlights the intricacies and effectiveness of each model in capturing detailed cellular characteristics. In Figure 3, we also present the capability of our proposed approach in generating cell image sequences against a variety of backgrounds given two reference background images. As shown, these backgrounds are precisely referenced from the reference background image used in the low-resolution video-to-video model, highlighting our method’s adaptability in replicating diverse cell environmental settings.

**Table 1:**
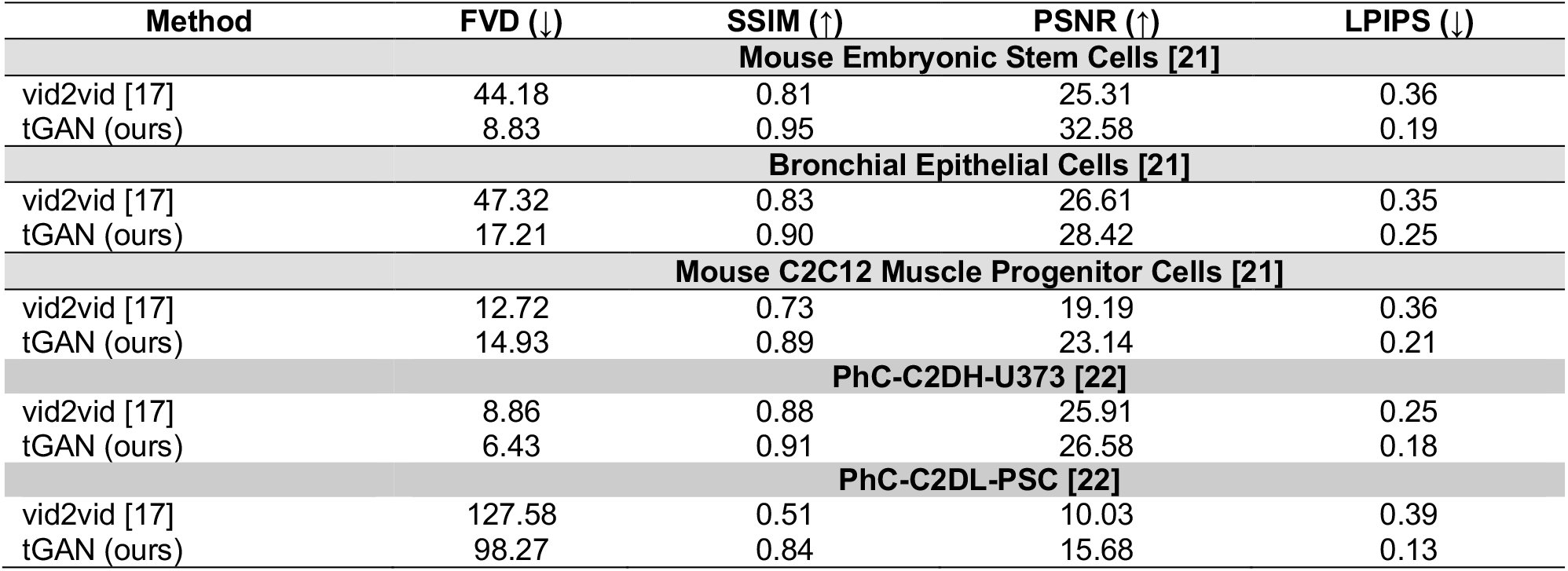
Comparative assessment of model performance. This table presents a comparison of our model’s performance across five different cell-type image sequences against the vid2vid model, measuring different video similarity metrics.

**Figure 2:**
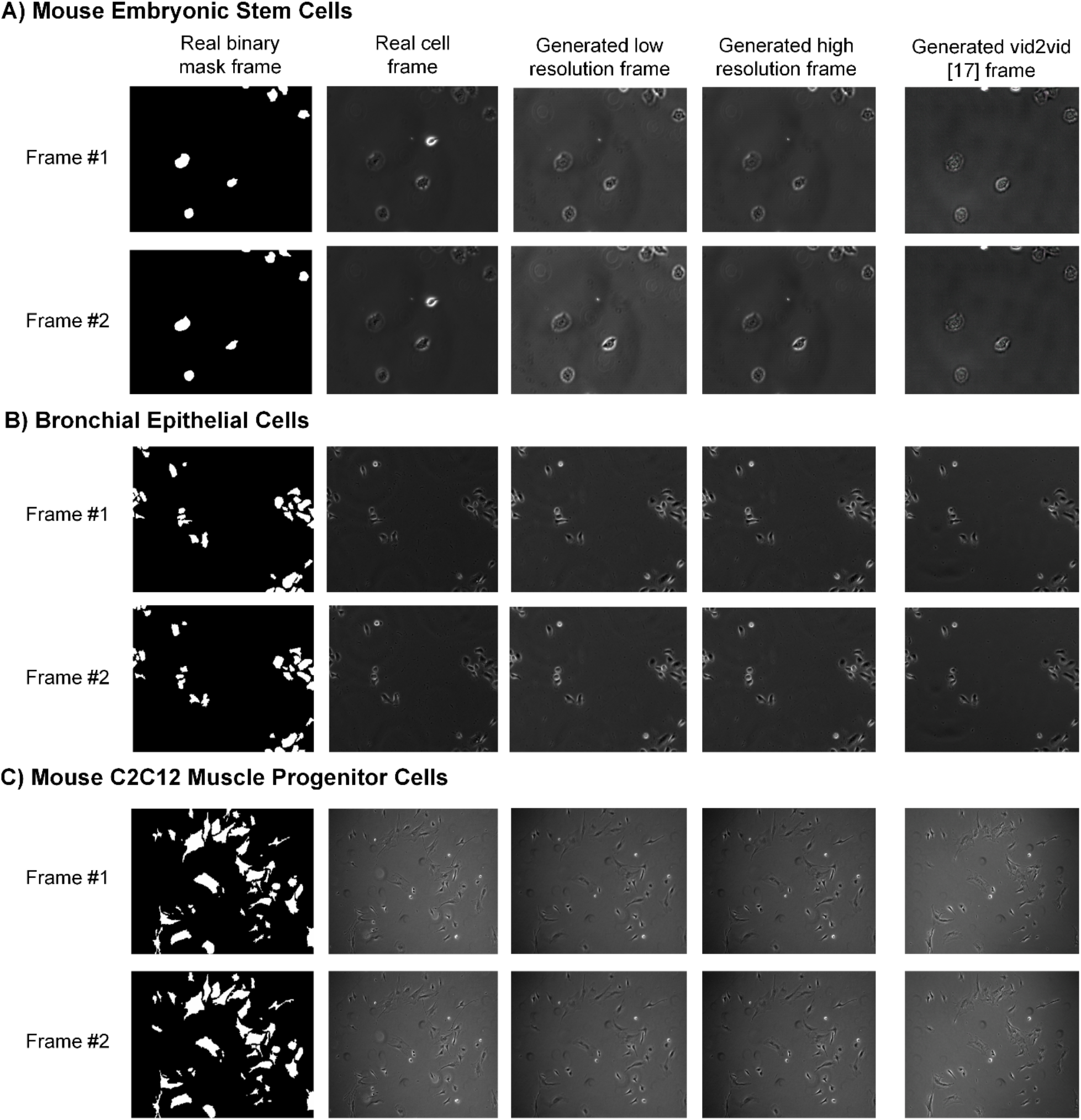
The examples of two consecutive synthetic cell images generated by our proposed tGAN model compared to the vid2vido model outputs.

**Figure 3:**
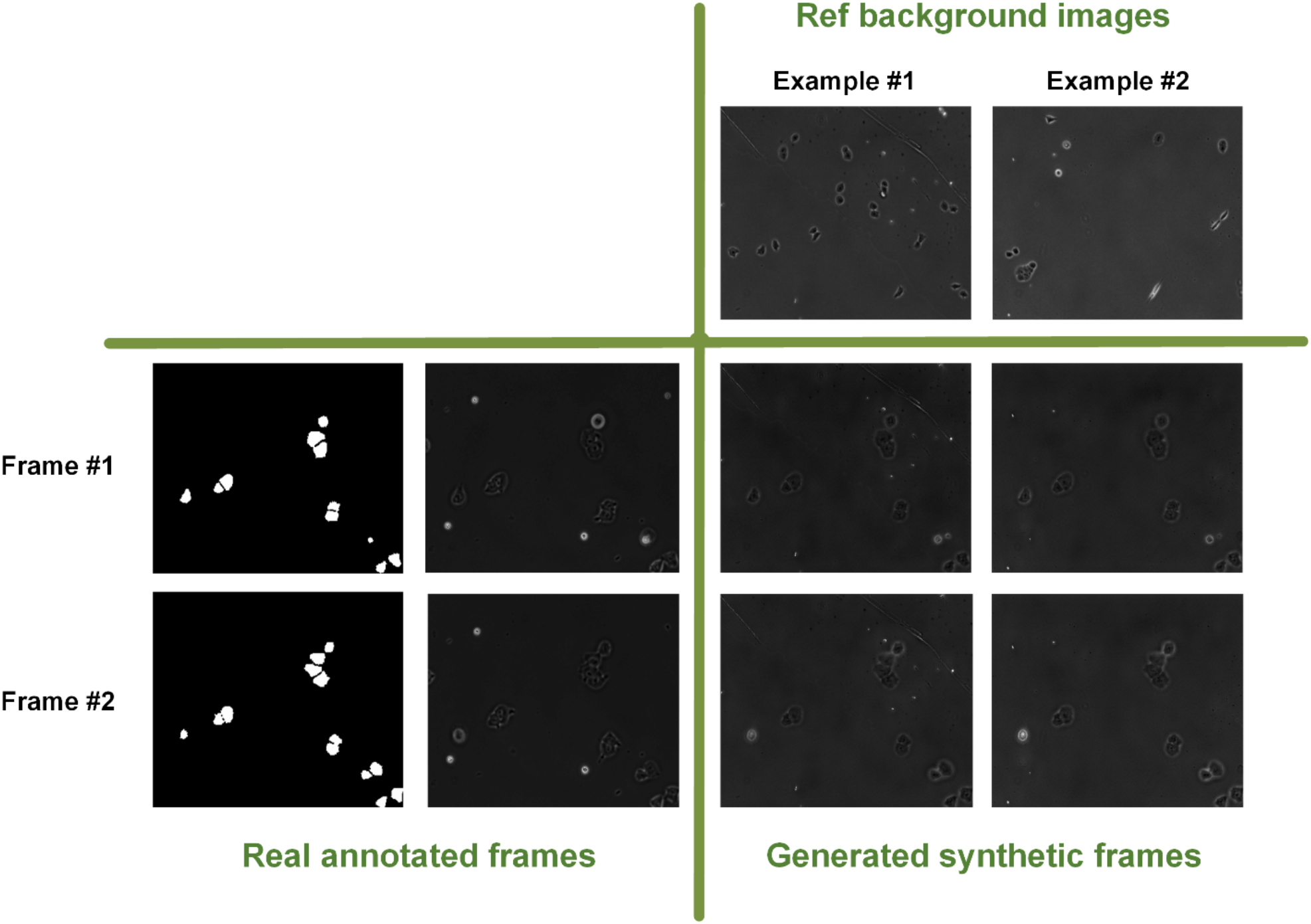
The examples of applying different reference background images for the same sample.

To test the stability of tGAN over time, we analyzed the FVD (Frechet Video Distance) score against the length of the tGAN-generated video in frames (Figure 4). We observed that the FVD scores exhibited only a slight fluctuation, approximately 1 FVD unit, when comparing videos ranging from 10 to 30 frames in length. This consistency in FVD scores, regardless of video length, underscores our tGAN model’s stability and reliability in generating high-quality video sequences over varying durations.

**Figure 4:**
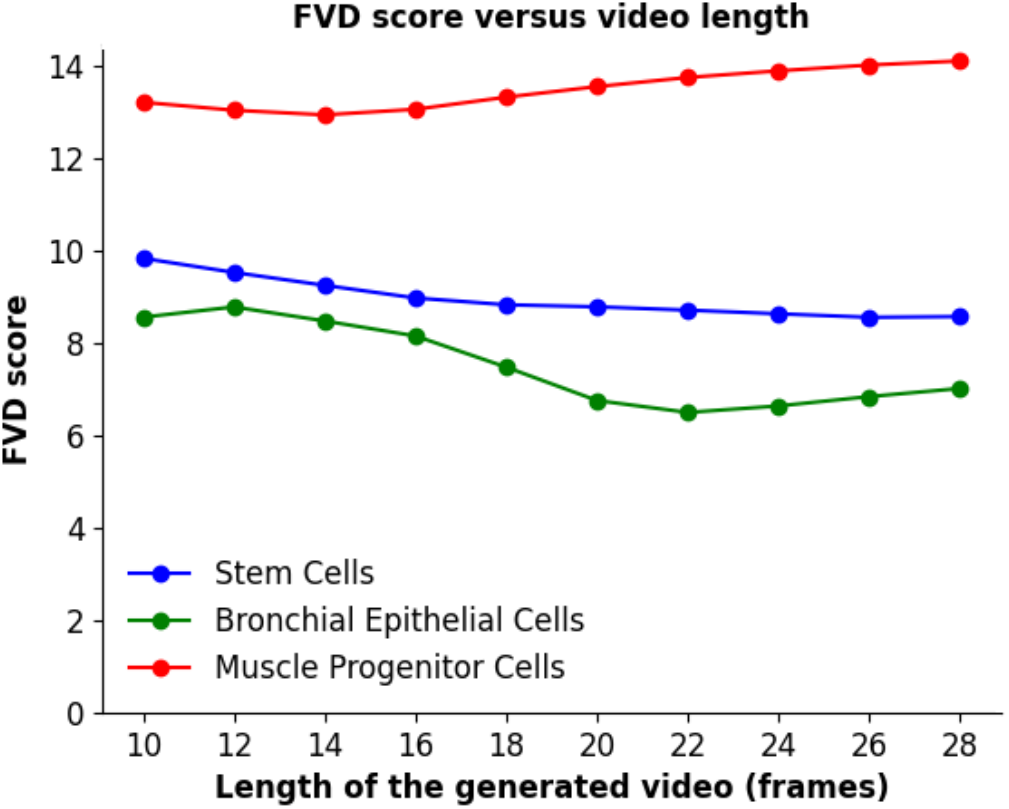
Frechet Video Distance (FVD) scores across different video lengths for three DeepSea cell type time-lapse video frames

We noticed that the tGAN training dataset (DeepSea and Cell Tracking Challenge Datasets) predominantly contained low and mid-density cell image sequences, which led us to investigate if our model could generate synthetic high-density cell videos of cells, a type not seen during training. This aspect is crucial as manual annotation of high-density microscopy videos is laborious and prone to errors. Therefore, we developed an algorithm specifically for creating synthetic high-density and colony-like time-lapse binary mask images, which is comparatively straightforward using conventional image processing techniques. These were then used as inputs for our tGAN generator during testing. We demonstrated that tGAN could successfully extrapolate from low and mid to high-density images, validating its ability to produce a broad spectrum of realistic cell images and highlighting its potential in various applications (Figure 5).

**Figure 5:**
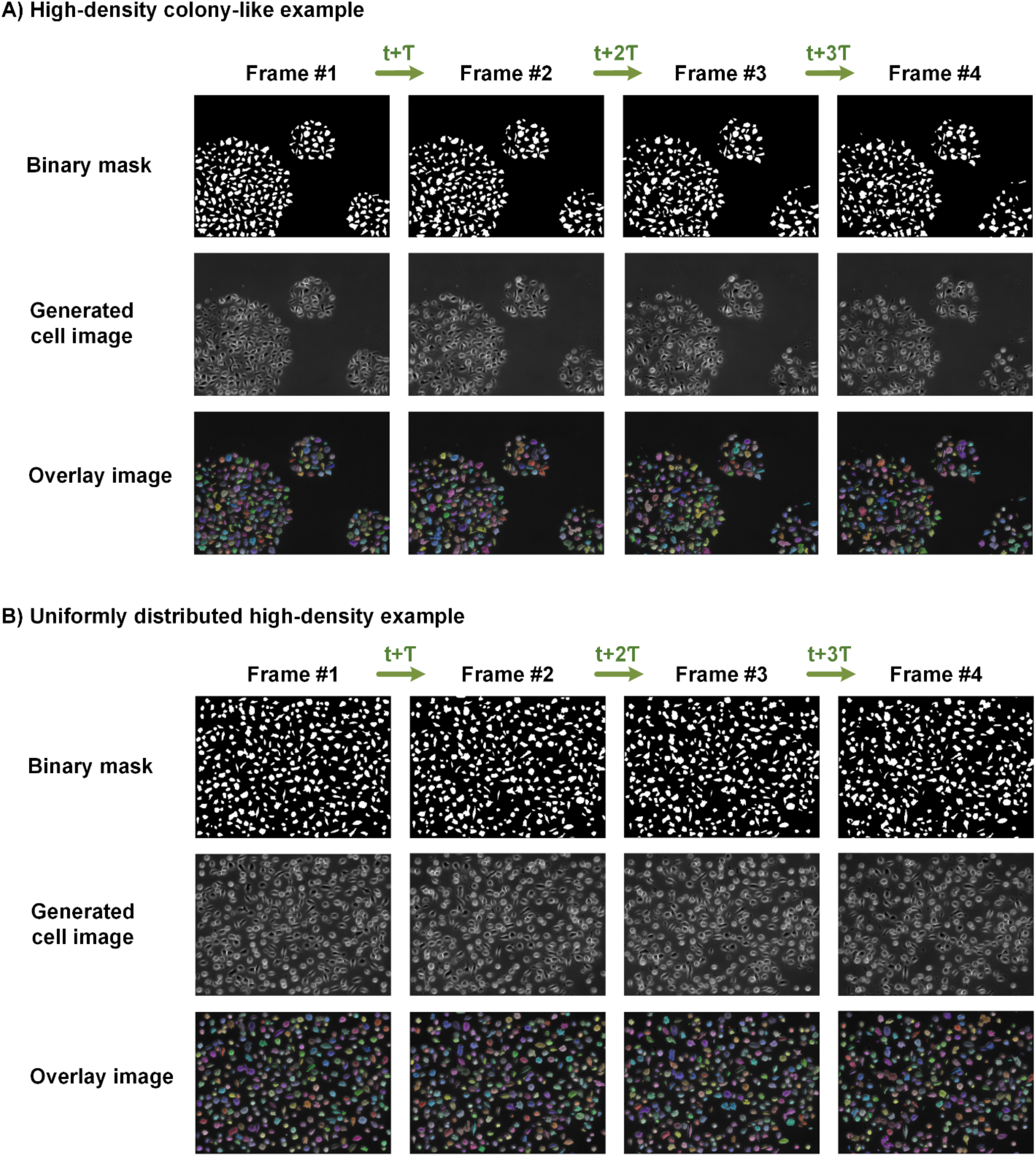
The examples of producing synthetic high-density and colony-like time-lapse cell video frames using our tGAN given synthetically generated binary mask images

Together, these findings demonstrate that tGAN can generate videos of cells with morphological details and temporal coherence similar to that of real videos of cells and across different annotated time-lapse microscopy datasets, further emphasizing the robustness and versatility of our approach.

### tGAN enhances performance of cell tracking models

Cell tracking remains a challenging part of bioimage analysis due to the dynamic nature of cellular behaviors, such as variations in cell shape, movement, and interactions within dense populations.

In addition, the scarcity of annotated data for training the sequence-based deep learning models, such as cell trackers, limits the application of deep learning models for microscopy analysis. We sought to test if the videos generated by tGAN can achieve the close cell tracking scores as real-time-lapse microscopy, thereby addressing the scarcity of annotated time-lapse video for training cell tracking modes.

This assessment is conducted using the DeepSea cell tracker [21] and Bayesian Tracker (btrack) [23], comparing the results to real annotated time-lapse videos from the DeepSea dataset. The evaluation leverages various object-tracking metrics that we described in our previous research [21], allowing for a detailed comparison of each cell-tracking model’s performance. This evaluation focuses on how closely our model’s tracking scores align with the ground truth annotated cell image sequences and align with our primary objective, which is to develop a GAN model capable of producing realistic annotated time-lapse microscopy images, addressing the scarcity of annotated data for training the sequence-based deep learning models, such as cell trackers.

As shown in Tables 2 and 3, the DeepSea tracker showed overall better results than btrack in all cases, likely because the model is already trained on the DeepSea dataset samples. More importantly, our tGAN model shows better and closer results to real annotated time-lapse microscopy cell images compared to the vid2vid model. This confirms that the video frame sequences generated by our tGAN model possess more realistic static and dynamic structures across frames, further validating the effectiveness of our approach in producing high-quality synthetic imagery.

**Table 2:** Quality evaluation of synthetically generated time-lapse microscopy sequences using DeepSea Cell tracking model, measuring different object-tracking metrics.

| Time-lapse test sequences | MOTA (↑) | MT (↑) | ML (↓) | Precision (↑) | Recall (↑) |
| --- | --- | --- | --- | --- | --- |
| **Mouse Embryonic Stem Cells [21]** |  |  |  |  |  |
| Real | 0.90 | 0.82 | 0.02 | 0.96 | 0.94 |
| Synthetic Vid2vid | 0.73 | 0.57 | 0.09 | 0.94 | 0.81 |
| Synthetic tGAN | 0.93 | 0.90 | 0.01 | 0.97 | 0.98 |
| **Bronchial Epithelial Cells [21]** |  |  |  |  |  |
| Real | 0.93 | 0.94 | 0.02 | 0.96 | 0.98 |
| Synthetic Vid2vid | 0.88 | 0.88 | 0.03 | 0.89 | 0.90 |
| Synthetic tGAN | 0.91 | 0.92 | 0.03 | 0.95 | 0.97 |
| **Mouse C2C12 Muscle Progenitor Cells [21]** |  |  |  |  |  |
| Real | 0.80 | 0.64 | 0.03 | 0.93 | 0.89 |
| Synthetic Vid2vid | 0.52 | 0.23 | 0.25 | 0.74 | 0.64 |
| Synthetic tGAN | 0.76 | 0.60 | 0.06 | 0.94 | 0.85 |

**Table 3:**
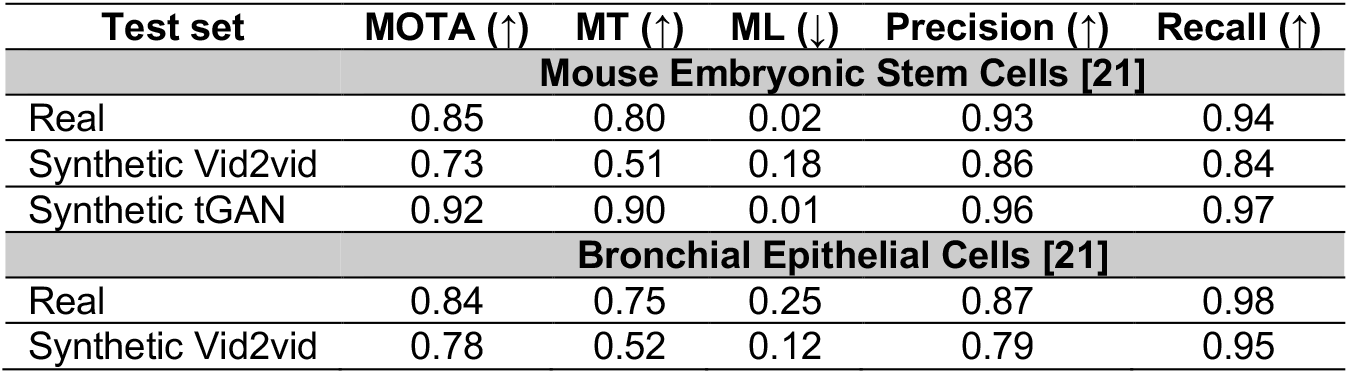
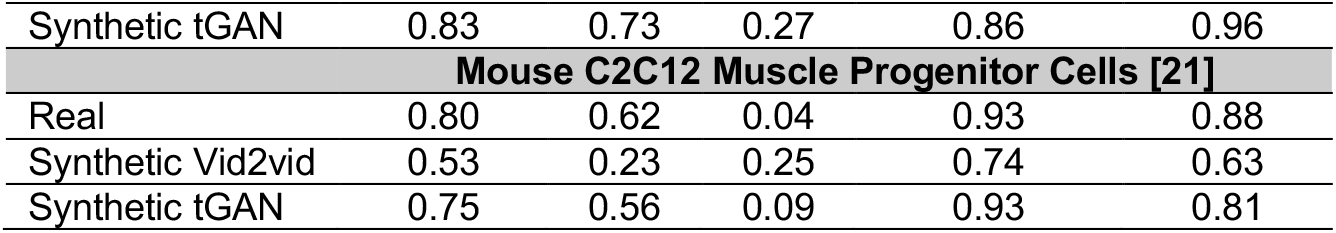
Quality evaluation of synthetically generated time-lapse microscopy sequences using btrack model, measuring different object-tracking metrics.

## Discussion

A foundational aspect of developing deep learning models, particularly in the field of biomedical imaging, is the availability of a large and diverse annotated dataset. However, the creation of such datasets for microscopy images is often hindered by the laborious and time-consuming nature of manual annotation. Addressing this bottleneck, our study introduced tGAN: a GAN-based super-resolution video generator designed to generate synthetic microscopy videos to address the scarcity of annotated live microscopy datasets, a field that demands high accuracy and detail in image processing. tGAN effectively circumvents the need for extensive manual annotation, generating realistic and diverse sets of synthetic annotated time-lapse cell images. Our study introduces a new approach to enhance the breadth of available training data for live microscopy to significantly boost the performance of segmentation and tracking models, particularly in scenarios with limited annotated datasets.

The model’s two-part structure adeptly handles both low-resolution and high-resolution image generation, a design choice that has proven valuable for tGAN to be computationally efficient. The integration of style and noise injection enhances the realism and morphological details of the images generated by tGAN. In addition, the inclusion of a reference background image ensures the adaptability of tGAN to diverse time-lapse scenarios with different background characteristics. We included different cell types in our study to demonstrate the performance of tGAN in a variety of challenging cell imaging scenarios. Our model’s superior performance in generating high-quality, realistic time-lapse microscopy videos, as evidenced by its outperformance of state-of-the-art models, underscores its effectiveness in handling complex microscopy images. This success is quantitatively supported by comprehensive metrics, which collectively affirm the model’s superiority in image quality, temporal coherence, and perceptual accuracy. Notably, our model’s ability to generate annotated time-lapse microscopy images to enhance the performance of tracking models marks a useful and novel advancement in the field. tGAN is also able to generate synthetic video cells with high density, which can address challenges associated with manual annotation of high-density cell videos for the development of segmentation and tracking models. By generating synthetic high-density cell binary masks and using them as the model inputs in the test phase, we demonstrated the model’s capacity to extend its application beyond the conditions experienced during training. This extrapolation is a practical benefit that can help reduce the time and effort required for manual annotation.

In conclusion, our tGAN model stands as a valuable tool with novel applications in microscopy imaging, capable of generating high-quality synthetic annotated time-lapse cell images across a spectrum of densities and cell types. While our current focus has been on two datasets, the DeepSea and Cell Tracking Challenge datasets, the model’s architecture can be extended to another dataset across different live imaging modalities. Future work should explore the model’s adaptability to even more diverse cell types, imaging conditions, and imaging modalities, as well as optimization of computational efficiency for larger datasets. The potential for further enhancing tGAN’s generalization capabilities remains an exciting prospect that we plan to pursue in the near future.

## Methods

### Dataset

In our study, we utilized two distinct training datasets from time-lapse microscopy, each representing different cell types, to ensure the robustness and generalizability of our proposed generative model across various biological contexts. These datasets are 1) Our recently published annotated dataset of phase-contrast images from the DeepSea collection [21], which includes a large set of accurately annotated phase-contrast time-lapse microscopy images of three cell types: Mouse Embryonic Stem Cells, Bronchial Epithelial Cells, and Mouse C2C12 Muscle Progenitor Cells. 2) The Cell Tracking Challenge dataset [22], a repository of 2D and 3D time-lapse sequences of fluorescent images featuring different cell types, such as PSC and U373 cells.

### Low-resolution video-to-video generative model overview

As illustrated in Figures 1B and S2, our low-resolution model is a sequence-based model inspired by the 2D-UNET architecture, known for its efficiency in image segmentation tasks [11]. This model is uniquely designed to process three types of input: the current consecutive mask images (e.g., two consecutive mask images), the previous generated synthetic cell image, and a consistent background image that serves as a reference for background visual features. The mask sequences act as the driving force, dictating the formation of the synthetic cell images, which are then superimposed onto the reference background. This approach allows for a more realistic and context-aware generation of cell images, which is essential for accurate subsequent analysis and ensures the consistency of the background visual features as well.

Central to our model’s architecture is the incorporation of specialized attention layers. These layers are strategically placed to merge features from three encoded inputs adaptively at different down-sampling stages. This process employs an attention mechanism, directing the model’s focus to the most pertinent features across the different inputs, thereby enhancing the detail and relevance of the generated images. In addition to structured feature integration, our model innovatively employs style and noise injection techniques in its decoding pathway, inspired by advancements in neural style transfer [24]. As shown in our previous research [12], this approach introduces a layer of variability and texture to the synthetic images, elevating their realism and authenticity. The style and noise elements are carefully modulated to complement the intrinsic features of the cell images, ensuring that the synthetic outputs are not only high in resolution but also rich in biological detail. Its ability to process and integrate multiple input types, coupled with the innovative use of attention mechanisms and style injections, sets a new benchmark in the field of synthetic image generation for time-lapse microscopy.

### Super-resolution image-to-image generative model overview

Following the generation of low-resolution synthetic cell image sequences, our approach employs a super-resolution generative model to further refine and enhance these synthetic images (Figure S3). This super-resolution model, similar to its low-resolution counterpart, is built on a UNET-based architecture [11] but with additional enhancements. This model similarly integrates style and noise injection techniques in its decoding path, which is influenced by StyleNet principles [24]. The style and noise injection significantly enhance the textural details and stylistic elements, contributing to the generation of more realistic and aesthetically consistent images, as demonstrated in our previous research []. The architecture features a sequence of encoder blocks that increase feature map depth, capturing intricate cell details. This is followed by a bottleneck process that prepares these features for nuanced reconstruction. In the decoder stages, the model combines upsampled features with style and noise information, progressively enhancing image resolution and quality. An additional upsampling layer in the decoder further increases the output resolution to 512×768, which can be adjusted to higher resolutions by modifying the model parameters.

### Discriminator architecture overview

The discriminator plays a pivotal role in distinguishing between real and generated images. We have chosen the PatchGAN architecture for the discriminator of both low-resolution and high-resolution training, enhanced with the addition of a linear attention layer in its early layers (Figure S4). This architecture and the inclusion of specific components are deliberate choices aimed at optimizing the discriminator’s performance. PatchGAN is known for its effectiveness in distinguishing fine details in images, making it a good choice for our purposes [13]. Unlike traditional discriminators that classify an entire image as real or fake, PatchGAN focuses on classifying smaller patches of the image. This approach is particularly beneficial for our model as it ensures that the generated images not only look realistic on a macro scale but also maintain high fidelity in finer details. In the measurement of our low-resolution discriminator loss, we employed a specific approach to enhance the model’s discriminative capability. At each training step, we concatenated the last n (e.g., two) real and synthetic (fake) frames along with their corresponding last two real binary mask frames. This concatenation provides the discriminator with a more comprehensive context, allowing it to assess not just the individual frames but also their temporal consistency and alignment with the binary masks. This technique is particularly effective in reinforcing the discriminator’s ability to discern subtle differences between real and generated image sequences, thereby sharpening the adversarial dynamic of the model.

The integration of a linear attention layer in the early layers of the discriminator is a beneficial approach. Attention mechanisms have gained popularity in various deep-learning applications for their ability to enhance model performance by focusing on relevant features while ignoring irrelevant ones [14]. The linear attention layer in our model allows the discriminator to prioritize certain aspects of the image, such as specific textures or patterns that are crucial for making accurate classifications. This focused approach improves the model’s efficiency and accuracy in distinguishing real images from synthetic ones.

### FlowNet architecture overview

In the training process of the low-resolution video-to-video generative model, we simultaneously trained a FlowNET model (Figure S5), a specialized component designed to calculate and integrate flow loss. This flow loss is crucial for accurately simulating the dynamics of cell movements, thereby enhancing the temporal consistency across video frames [15]. We incorporated this additional loss metric to optimize the generative model’s weight updates, specifically aiming to maintain temporal coherence in the synthesized video sequences. This strategy ensures that the generated sequences not only mirror real-world temporal dynamics but also enhance the realism and scientific applicability of the generated images.

The optical flow loss, calculated by our FlowNET, is pivotal in maintaining the integrity of temporal dynamics across generated frames, aligning these dynamics closely with real-world observations to enhance the realism and scientific utility of the synthetic images. It’s important to note, however, that the application of flow loss is selectively applied; it is not utilized in the high-resolution model, which focuses primarily on enhancing image detail through image-to-image translation, thus reducing the need for temporal consistency in that specific context.

### Augmentation process in training

We employed a robust augmentation pipeline to enhance the training of both the generators and discriminators. This process involves applying a series of video-level augmentation functions to each training sample, designed to introduce variability and improve the model’s generalization capabilities. As illustrated in Figure S6A, we applied some mostly used conventional image augmentation functions, including random adjustments in histogram equalization, sharpness, brightness, and contrast, as well as horizontal and vertical flips, cropping, saturation modifications, and the addition of Gaussian noise and blur. The training algorithm executes a sequence of the provided augmentation functions for each cell and mask video pair with a pre-defined probability value ‘p_vanilla’. In the requested augmentation pipeline, each function is randomly chosen with a consistent probability of 50% and is also applied in a randomized sequence.

In the training of GAN models, the limitations of conventional augmentation become apparent, particularly in its inability to significantly diversify the generated images when the training dataset size is limited. To address this challenge, the concept of differentiable augmentation, detailed in [16], proves to be invaluable. This approach has also been validated in our previous research [12]. Differentiable augmentation applies the same random augmentations to both real and fake samples in a way that is differentiable with respect to the model parameters. This approach encourages the discriminator to mitigate overfitting and improve training stability, making it particularly beneficial for GANs trained with limited data, thus causing the generator to produce more diverse images, thereby improving the overall image generation performance. In the training process, we ran five distinct differentiable video-level augmentation functions, such as random contrast, brightness, cutout, translation, and saturation (Figure S6B). The application of each augmentation is controlled by a predetermined probability variable, ‘p_diff’. To promote unbiased representation and randomness in the training data, these augmentations are applied in a random sequence. Each function has an equal chance of being selected, set at a 50% probability.

### Loss functions and their rationale

We employed a combination of other critical loss functions (Equations 1-3) to guide the models effectively during the training process. These include perceptual (VGG) loss [25], L1 loss, and discriminator loss, each serving a specific purpose and contributing to the overall performance and accuracy of the model. The perceptual (VGG) loss and the L1 loss, respectively, ensure perceptual and pixel-wise similarity between the generated and real images. These losses focus on high-level features and granular accuracy. The discriminator loss, Mean Squared Error (MSE), plays a dual role: for the generator, it gauges effectiveness in deceiving the discriminator, and for the discriminator, it evaluates its capacity to differentiate real from synthetic images. This adversarial loss is key in driving the generative process toward producing images that closely mimic real ones.

We also, in the training process, assigned a specific weight (w1-w3) for each of these loss functions aiming to ensure a balanced contribution during the optimization process. This weighting is crucial as it fine-tunes the impact of each loss function according to its relevance and importance in the image generation task. It is important to note that we do not employ flow loss in the training process of the super-resolution model, as the FlowNET, which calculates flow loss, is not utilized in this phase of the training. This decision is based on the super-resolution model’s focus on image-to-image translation rather than temporal video sequence generation.

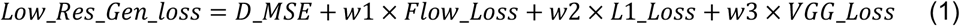

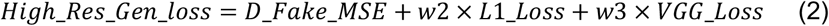

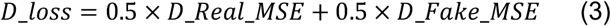

## Code availability

The Python scripts that implement the methodologies we developed are publicly available for download at our GitHub repository: https://github.com/abzargar/tGAN. Additionally, the image datasets used in our study can be accessed through a link on the repository page, enabling straightforward replication and further research.

## Author Contribution

A.Z.K and S.A.S were pivotal in conceptualizing the project and played a significant role in manuscript preparation. Both A.Z.K and N.M developed the related Python scripts and prepared the manuscript materials.

## Acknowledgments

We would like to thank the members of the Shariati lab for their feedback on the manuscript. We also would like to thank Ryan Shariati for his critical discussion on the potential application of synthetic live microscopy images.

---

**Figure S1:**
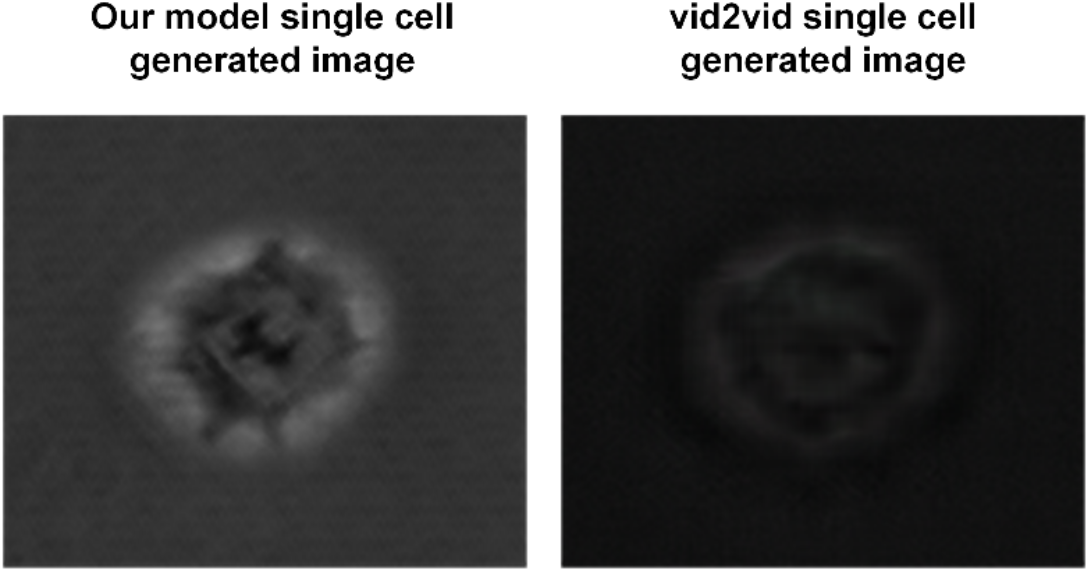
Comparative performance of our model vs. vid2vid in replicating fine visual features of single stem cells.

**Figure S2:**
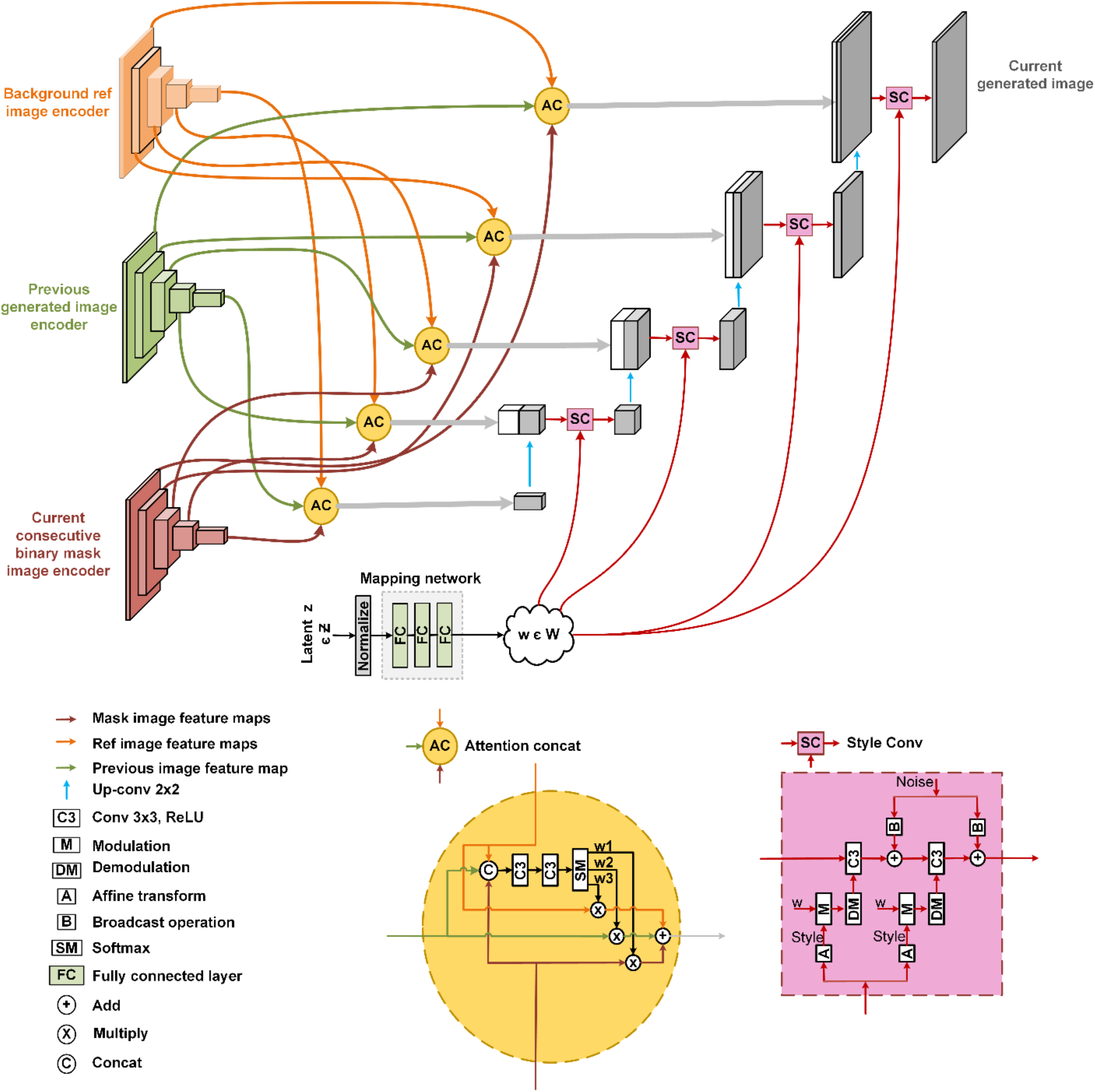
Detailed architecture of the low-resolution sequence-based generative model utilizing 2D-UNET encoding and decoding pathways, attention mechanism, and style injections.

**Figure S3:**
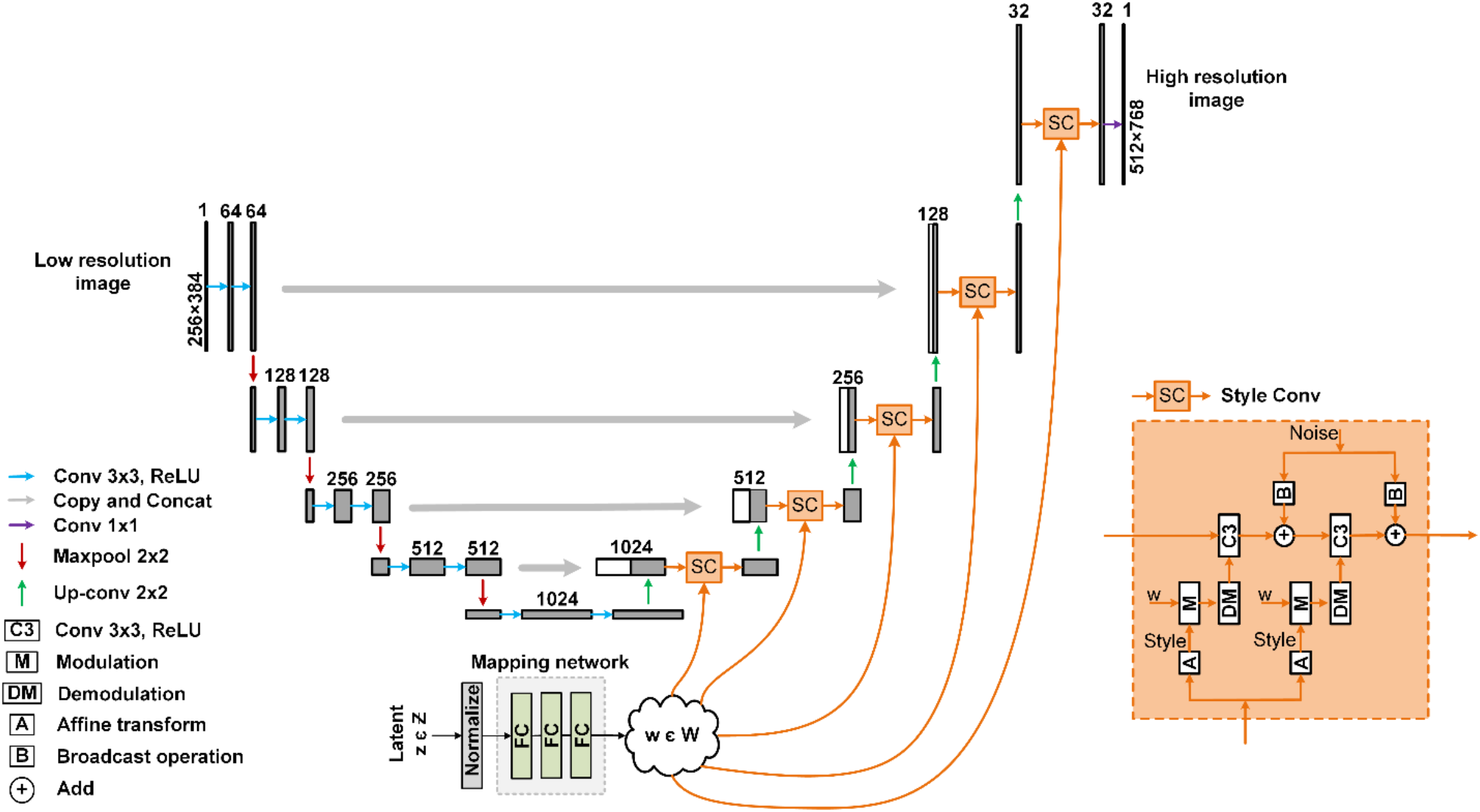
Super-resolution generative model architecture for refining low-resolution synthetic cell image sequences, integrating UNET-based design with style and noise injection techniques.

**Figure S4:**
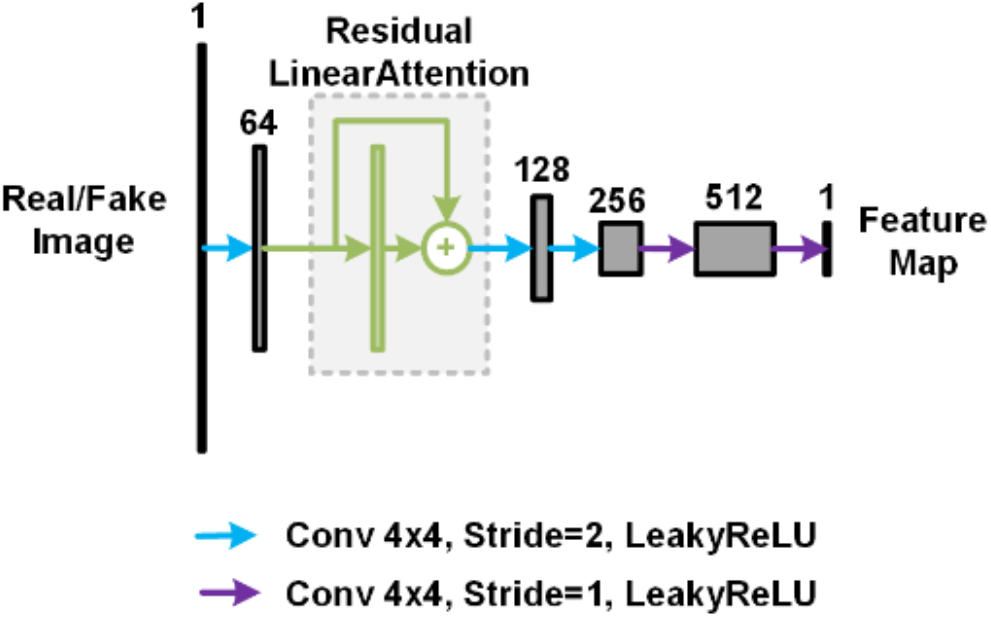
Discrimination model architecture employing PatchGAN techniques with linear attention.

**Figure S5:**
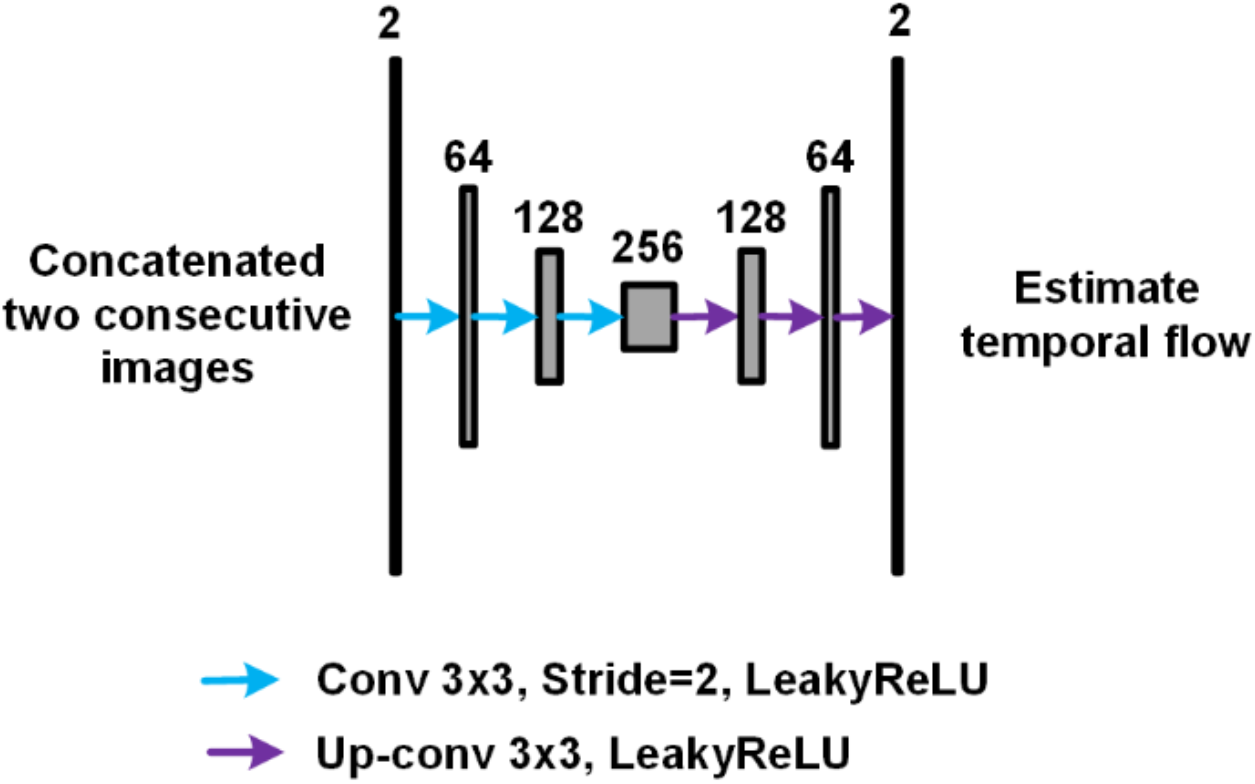
Our FlowNET model architecture used in the training process of the video-to-video low-resolution generative model

**Figure S6:**
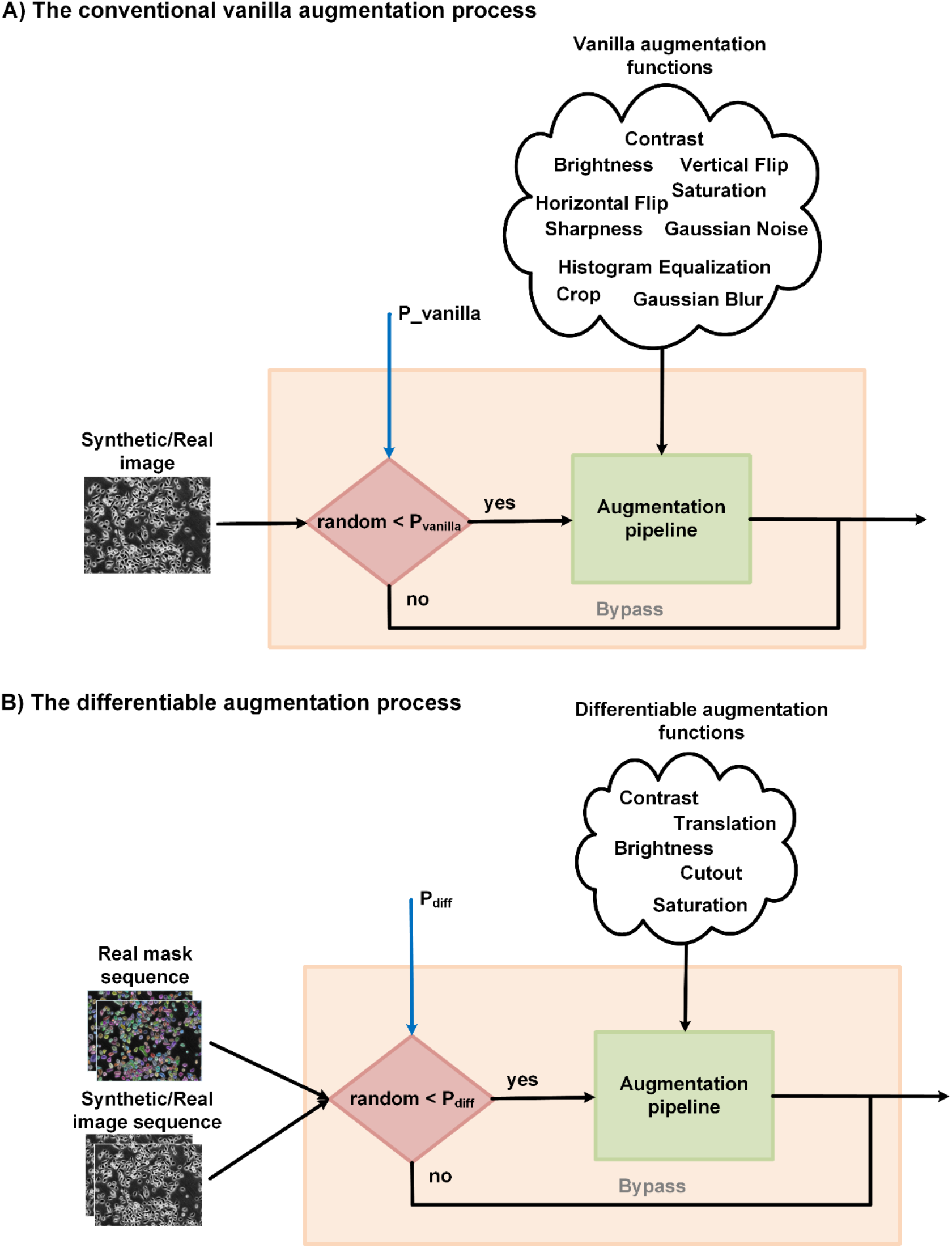
The proposed video-level augmentation approach used in the training process of our generative model.

